# Anatolution, an online platform for consensus morphology

**DOI:** 10.64898/2026.02.16.706144

**Authors:** Daniel J. Miller, Blaise Gratton, Zoe LeBlanc, Jon H. Kaas

## Abstract

**Introduction:** Supervised statistical learning for cell-level segmentation and morphometry in optical microscopy is limited less by algorithmic capacity than by the scarcity of reliable, expert-validated ground truth. In comparative neuroscience and quantitative histology, where classical stains such as Nissl’s method remain the primary means to study cellular morphology, this bottleneck is acute: manual annotation is expensive, subject to individual bias, and rarely performed at the scale or consistency that computational approaches demand. No existing platform integrates a stain-specific bioimage segmentation protocol, a structured multi-annotator workflow, and consensus-based quality control into a single pipeline from image ingestion to machine-readable training data.

**Methods:** We present Anatolution, an open-source, web-based platform designed to address the gap of quality annotations at https://anatolution.herokuapp.com/public-tool/. Anatolution organizes microscopy images, including 2D arrays or 3D volumes, into project workspaces where multiple annotators independently label cellular structures against a shared computer vision catalogue. This design enables systematic inter-rater and intra-rater reliability assessment, with consensus derived from agreement across annotators rather than from any single expert’s judgment. The platform enables the export of aggregated labels or annotation datasets for downstream statistical learning methods. We describe the system’s architecture, its Nissl-specific segmentation pipeline, the consensus annotation workflow, and validation of inter-rater reliability.

**Conclusion:** Across 20+ histological annotation containers annotated by up to 15 independent raters, consensus boundary agreement increased monotonically with annotator count, reaching a median Dice of 0.79 against the full-rater reference at seven annotators, with top-tier containers achieving leave-one-out ceiling values of 0.621–0.769 for cell-body segmentation. The segmentation pipeline provided effective spatial anchoring, with 88% of consensus-annotated polygons containing at least one algorithmically detected seed. Anatolution provides open-source infrastructure for producing consensus-validated training data from classical histological preparations, addressing the primary bottleneck limiting supervised learning for cell-level morphometry.

## Introduction

Quantitative neuroanatomy begins with the premise that understanding brain function requires accurate measurement of cellular organization and microstructure (1,2). For decades, the field has sought robust methods to quantify the number, distribution, and morphology of cells and fibers, and to relate these measurements to higher-order anatomy and behavior (3). Yet persistent challenges in achieving consistency across individuals, laboratories, and protocols continue to limit what can be reliably inferred about brain organization (4–7). Unfortunately, these challenges are not merely technical but interpretive, and define the boundaries of what sparse, noisy, and difficult-to-reproduce measurements can reveal (8). Basic, ubiquitous technical artefact remains a challenge in particular for interdisciplinary fields like comparative neuroscience that rely heavily upon bringing together observations across taxa that are often subject to unavoidable differences in post mortem sample processing (3). Coincidental to the limit this technical challenge presents is an opportunity for the field to unite in the analysis of a single stain, like Nissl’s method (9,10), that is convenient, affordable, and robust to variation in tissue processing methods (9), as opposed to more challenging molecular methods (11–13). The fundamental task of identifying, classifying, and counting cells in stained tissue sections sounds simple but has proven remarkably difficult to automate (14–16) at the accuracy levels that rigorous quantitative inference demands.

The multiscale integration of measurement in neuroscience depends critically on expert-validated annotations of the ground truth neurobiology of interest to inform basic and medical applications. The growing use of noninvasive neuroimaging increases demand for quantitative links between microscopic tissue features and imaging-derived metrics, particularly in translational contexts where biomarkers are used to infer cellular architecture. Reliable annotation datasets can support co-registration, quantitative comparisons, and model calibration, strengthening the interpretability of imaging biomarkers across species and conditions (14,16,17). Improving annotation infrastructure is therefore not only a computational requirement but a foundational step toward unifying measurement across scales in neurobiology to illustrate first principles of brain function, and ultimately evolution.

Stereological methods emerged as the gold standard for quantitative histology beginning in the 1960s, when inconsistencies across clinical laboratories prompted Hans Elias to organize the first International Society for Stereology (8). These methods provide a framework for statistically principled sampling of three-dimensional morphological features from two-dimensional sections (8,14). Unfortunately, the high cost and steep expertise requirements alongside the labor-intensive implementation of these methods has limited widespread adoption, particularly for studies requiring large sample sizes, extensive anatomical regions, or repeated measurements across conditions (3,8). Alternative approaches such as the isotropic fractionator offer faster throughput but sacrifice spatial information by homogenizing tissue into nuclear suspensions (3,9,18). The gap between what is theoretically measurable and what is routinely measured remains substantial (14), motivating approaches that can preserve statistical rigor while increasing scale.

Recent advances in computational capacity and statistical modeling have raised the possibility of high-throughput quantitative histology (19). Deep learning approaches can now classify whole-slide images with complex morphologies (17), extract high-resolution features (19), and support analyses at scales previously impractical (9,10,20,21). In principle, emerging single-instance segmentation methods can bridge microscale to mesoscale to macroscale structural datasets, but the performance of these methods remains constrained by heterogeneous tissue, variable staining, subtle morphological classes, and demanding accuracy targets (15,22). As segmentation targets approach near-perfect accuracy — necessary when errors accumulate across millions of objects — the demands on training data grow (16,23). The primary bottleneck is thus not the lack of algorithms or computational depth, but the scarcity of sufficiently large, accurate, and biologically meaningful training and evaluation datasets (14,15,17,19,23–25). Contemporary reviews consistently identify training data quality as the central constraint on deep learning performance in histological image analysis, with supervised methods outperforming unsupervised approaches when adequate annotations are available (14,15,26). Many authors note that manual annotations are “prohibitively” expensive, yet this leaves the problem unsolved and thus we propose to try making annotation more efficient and reliable in pursuit of a quantitative cellular histology (14,26).

High-quality ground truth is expensive because it requires domain expertise, careful definitions of what constitutes an object, and handling of artefacts and ambiguous morphology (24,25,27,28). Existing tools — CellProfiler (29,30) for modular image analysis pipelines, Ilastik for interactive machine learning segmentation (31), and Cellpose (32) for deep learning-based cell detection — provide powerful infrastructure but are not designed to address the annotation problem itself as a problem of rater reliability. Tools to label data in images abound, but frameworks designed to explicitly manage rater agreement during curation are not yet standard, despite calls for higher quality standards. The single biggest source of variance in morphometric datasets is often the investigator herself, which demands redundancies to estimate and reduce individual bias (3,14,23). Even a relatively low degree of ambiguity among skilled investigators — perhaps ten percent disagreement on individual objects — compounds catastrophically when scaling to millions of measurements, as different raters disagree about different parts of the decision space. Accordingly, improving annotation quality is not a peripheral task, but a central prerequisite for advance.

The mismatch between human visual competence and current computer vision performance on routine but critical measurement tasks in neuroanatomy remains an underappreciated dimension of current bottlenecks to quantitative anatomy (16,25,33). Primate visual evolution yields high-acuity pattern recognition that allows experts to integrate context, discount artifact, and interpret ambiguous boundaries with a flexibility that automated systems cannot yet match (34). Conversely, humans are poorly suited to exhaustive counting and repetitive measurement at scale, whereas computers excel at high-volume standardized operations given that the task has been formalized. This complementarity suggests an approach that does not attempt to replace human expertise but operationalizes it — using human judgment where it is essential and computation where it is advantageous — to produce reliable, scalable morphometric datasets. The field of computer vision is increasingly recognizing the need to augment human expertise through “human-in-the-loop” strategies, particularly for specialized domains where transfer learning from generic datasets performs poorly (14,23,25,32).

Consensus-driven annotation provides a principled and general way to increase machine training data quality. Specifically, agreement between trained experts reduces low-level errors and stabilizes definitions of morphological objects for downstream computation. When multiple trained observers contribute annotations and disagreements are detected, reviewed, and resolved prior to training, the resulting labels can better approximate a robust interpretation of morphology than any single rater’s annotations alone. Such consensus processes are especially valuable when the ground truth is intrinsically uncertain due to taxonomic variability, staining quality, tissue damage, partial volume effects, or ambiguous cell boundaries (15,23,24). Importantly, rater reliability metrics developed in the stereological literature — including intraclass correlation coefficients for assessing agreement — can serve simultaneously as quality control measures for training progress and as indicators of dataset reliability (3,9,14,19,23,35). In effect, consensus transforms annotation from purely manual labor into a structured scientific procedure that explicitly manages uncertainty and improves reliability as dataset size and annotator count grow.

Here we introduce Anatolution, an open-source, web-based platform designed to make consensus annotation practical for the comparative neuroscience community. Anatolution bundles three components into a toolkit: (1) a segmentation pipeline tailored to Nissl-stained cells, (2) a browser-based annotation interface supporting simultaneous independent annotation by multiple users, and (3) a workflow architecture organized around consensus with integrated agreement protocols and quality control reliability metrics. Computationally derived cell seeds serve as an objective backdrop for annotation, providing a shared reference frame to integrate inter-annotator comparison in the case of doubt as freehand annotations vary across users and require calibration. The platform supports configurable labeling vocabularies, structured intra-rater and inter-rater reliability assessment, and exports standardized, machine-readable datasets. In this manuscript, we describe Anatolution’s workflow, system architecture, and validation procedures, and present evidence that the consensus-first approach produces reliable, reproducible annotations whose quality is measurable and improves with annotator count — establishing the infrastructure through which consensus-derived training data can systematically improve downstream model performance. The problem of annotation reliability is perhaps most acute for evolutionary neuroscience, where the lens of comparative neuroanatomy has revealed extraordinary taxonomic diversity. We thus introduce Anatolution as part of this special issue on mapping neurobiological diversity as a facilitative toolkit for the intrepid evolutionary scientist.

## Methods

### 2.1 Philosophy and Framework

Anatolution is designed as research infrastructure for computational scientists who engage programmatically with their data and who will integrate annotation outputs into their own analysis pipelines. The platform consists of a graphical user interface (GUI) at https://anatolution.herokuapp.com/public-tool/ where users can upload images and generate custom annotations in a variety of file formats for export and use with computational modeling. Unlike general-purpose tools that prioritize accessibility for non-technical users, the current method prioritizes flexibility, transparency, and direct access to underlying data structures for our community of users who can train a segmentation model, write a SQL query, and parse a structured data export. This orientation reflects the reality that high-quality morphometric training data ultimately serves downstream statistical learning workflows that require technical expertise to implement (27,36). Accordingly, platform functions are built for efficiency—particularly data export—and are currently implemented through graphical interface to quickly acquire label data (27,29). Users have a basic tabular access to their variably annotated data, including per-user and per-project isolation, spatial coordinates, marker vocabularies, and three-dimensional contour geometries. The public tool is designed to facilitate the generation of machine-readable computer vision datasets and foster the development of an active deep learning ecosystem where members of the community may join or lead collaborative consensus projects.

### 2.2 Platform architecture and data management

Anatolution operates as a platform for unauthenticated users to generate datasets on a case-by-case basis via the public tool at https://anatolution.herokuapp.com/public-tool/. Each project defines a configurable vocabulary of annotation markers (e.g., cell-type labels, quality-control flags) and a set of images or image volumes to be annotated. Image sets may represent serial sections through a three-dimensional volume along the Z-axis, where physical dimensions and Z-spacing are recorded by the user as project metadata, or 2D arrays as in whole-slide image (WSI) stitching. A given image set can be assigned to multiple projects, with annotations stored separately per project, enabling independent analysis of the same tissue under different annotation protocols or by different user groups. All annotations are recorded per-user, preserving the independent observations required for subsequent consensus analysis and rater reliability assessment (37,38). The images are first uploaded and then organized into project-specific groupings, often corresponding to serial sections through a three-dimensional volume or adjacent tiles within a 2D stitch or whole-slide image. Annotators navigate through image groups along the determined dimension (Z or XY), marking or tracing objects of interest using configurable annotation markers (e.g., neurons, astrocytes, or regions of interest). Completed annotations are immediately available for download and can be exported in standardized formats for downstream computational pipelines (see Section 2.6).

### 2.3 Annotation interface

Anatolution provides a browser-based annotation interface for machine-readable bioimage dataset curation. The interface is designed around two core requirements: (i) enabling precise morphological annotation of cellular structures, including three-dimensional extent, and (ii) supporting independent multi-annotator workflows required for statistically grounded consensus. Annotation is organized around probes—conventionally volumetric image sets typically comprising 13–17 focal planes acquired at ~1 µm axial spacing (approximately 15 µm total depth in this report), or stitched tiles from a WSI. Each probe is subdivided into columns or rectangular fields of view (~20–25 µm lateral extent in this report) that serve as bookkeeping units for systematic tissue coverage and provide a natural reference scale for cellular structures. Within each column, annotators delineate objects by placing polygon vertices directly on the image. Vertices are added sequentially to form a closed boundary and can be repositioned or removed through standard interactions. Each polygon is stored as an ordered coordinate sequence, with vertex positions recorded as percentages of image dimensions to ensure resolution independence across display contexts and exports. For three-dimensional annotation, users navigate the Z-stack using a slice interface (keyboard shortcuts can be enabled for rapid traversal), placing polygons on one or more focal planes as needed. Polygon boundaries are stored per plane and linked by object identifiers, enabling volumetric reconstruction of each annotated structure across depth. Annotators may optionally toggle overlay visibility to inspect the raw image without visual interference from existing markings. Object categories are defined at the project level through configurable marker vocabularies, allowing each project to tailor annotation schema to staining method, tissue type, and scientific aims. For example, Nissl-stained workflows often use cell-type markers (e.g., neuron, astrocyte, oligodendrocyte, microglia, endothelial) together with quality-control markers such as artefact (e.g., incomplete or damaged objects, out-of-plane profiles, or non-cellular structures) and unknown for cases of low classification confidence. The unknown label is an explicit design feature that records uncertainty rather than forcing a potentially spurious decision as this uncertainty can then be quantified across raters and prioritized during later consensus deliberation. Critically, annotators working on the same project annotate identical image sets independently and are not visible to other users during the annotation phase. This separation is structurally necessary for the consensus workflow, ensuring that agreement reflects genuine convergence of judgment rather than anchoring or social conformity. When computational seed detections or other algorithmic overlays are available, they provide a shared visual reference that standardizes what annotators evaluate (i.e., candidate objects and coverage) without dictating the morphological label assigned.

### 2.4 Consensus Rater Reliability

The central methodological contribution we report is a consensus-first annotation strategy designed to address the persistent challenge of rater reliability in microstructural cell morphometrics (3,9,10,21). Rather than treating annotation as an individual task whose outputs are later compared for quality control, Anatolution structures annotation as a collaborative process in which multiple trained observers independently label identical image sets through the graphic user interface (GUI), and consensus is derived from convergence (39). Computer vision is leveraged primarily for recordkeeping to ensure the systematic coverage and tracking of candidate objects, while biological classification and boundary delineation remain the province of trained human judgment.

Scholars new to the annotation process undergo structured training prior to participation in our record-heavy consensus data collection method. Training begins with directed study of morphological criteria for cell-type identification in Nissl-stained tissue (3,9,39) to disambiguate neurons from glia and endothelial cells based on soma size and shape, staining intensity, nuclear features, and nucleolar morphology. Trainees compile a personal reference guide and then complete supervised practice annotations within Anatolution. The training period spans approximately one semester (roughly 16 weeks), during which each trainee typically spends 10 hours per week collecting data, with approximately 3 hours of structured mentorship weekly to calibrate classifications and boundary placement. Annotators learn the workflow by maintaining detailed notes of consensus agreement as a record for later troubleshooting and refinement. Annotators are considered *intermediate* after one semester (~48 hours consensus) and *advanced* after two or more semesters (~150 hours consensus), though the reliability data generated by the workflow serve as the ultimate determinant of annotator quality.

Prior to probe annotation, groups of four to five annotators convene in a facilitated consensus session led by a senior scientist who is typically also the project administrator. Scrolling through the images from the assigned probe, the group collectively examines candidate objects, discusses morphological features relevant to classification, and reaches agreement on object identity and the focal plane(s) in which each object is visible. The group documents its decisions—including objects whose identity is uncertain and the basis for that uncertainty—in a shared reference document accompanied by screenshots. This deliberation establishes shared criteria for the probe, provides a record available for mentor review, and surfaces ambiguous instances that later inform interpretation of inter-rater variability. The project-level marker vocabulary used for labeling (cell-type and quality-control markers) is fixed for the project, and annotators may assign an explicit *unknown* label when classification confidence is low, thereby recording uncertainty rather than forcing a potentially spurious decision.

Following deliberation, each annotator independently annotates the identical probe volume within Anatolution on their own workstation browser. Each annotator traces polygon boundaries for every object assigned for annotation, working systematically through columns and focal planes. At the midline plane—defined operationally as the focal plane in which the soma cross-section is largest and the boundary is most sharply resolved—annotators trace the full object boundary. In planes above and below the midline, where boundaries may be partially out of focus, annotators trace only the contour that remains optically resolved, using the transition between focused and defocused tissue across the Z dimension as a guide. During independent annotation, each annotator’s polygons are stored separately and are not visible to other annotators. This separation is structurally necessary as deliberation establishes *what* each object is and *where* it is located, while independent annotation captures *how* each annotator delineates the boundary, ensuring that agreement reflects genuine convergence of judgment rather than anchoring to a shared tracing.

Here we report the use of computer vision methods as visual bookkeeping and structured biological topologies to guide morphology and augment user domain expertise. Computationally derived candidate seed locations (described in Section 2.5) are displayed as a shared visual reference to support bookkeeping and structured coverage assessment. Seeds provide a completeness check by logging and flagging even false-positive detections for review, enabling targeted adjudication of omissions and systematic accounting of coverage (8,35). The computational backstop comparison provides a cross-check that neither source alone could offer—seed detections promote completeness, while human consensus promotes biological validity, including in the case of real morphological ambiguity (9).

Consensus is computed by comparing annotations across users at the object (or pixel) level(s). For each annotated object, a centroid is first computed from each user’s polygon (per plane for 2D contours, or an average or weighted centroid across planes for 3D objects). Centroid proximity may be used to establish correspondence among polygons referring to the same structure. When multiple candidate matches occur (e.g., split/merge disagreements), correspondence is resolved using maximal spatial overlap (area/volume intersection-over-union) as a tie-breaker, and unresolved or ambiguous cases are flagged for manual review. Agreement is quantified as the overlap among corresponding polygons, computed as area overlap for 2D boundaries and volume overlap for 3D reconstructions. When annotators demonstrate comparable reliability on the relevant marker classes, consensus boundaries are derived by majority vote—pixel-wise inclusion across raters.

Where applicable, Simultaneous Truth and Performance Level Estimation (STAPLE) provides a probabilistic consensus mask from multiple segmentations (14,38). When annotator skill varies, as assessed by leave-one-out Dice coefficients computed from pairwise annotation comparisons, individual contributions are weighted prior to aggregation, with weights bounded to the interval [0.5, 1.5] and normalized across annotators within each consensus set. This bounding ensures that high-agreement raters exert proportionally greater influence while maintaining a floor that prevents any rater’s contribution from being discounted entirely.

Rater reliability is treated as a continuous quality control measure throughout the annotation lifecycle (38). Inter-rater agreement (across annotators) and intra-rater agreement (within annotators on repeated material) are computed as annotations accumulate, functioning both as indicators of training progress and as quality measures for the resulting dataset (8,35,37,38). These metrics enable project administrators to identify annotators requiring additional training or recalibration before their annotations are incorporated into consensus and to prioritize ambiguous cases (including *unknown*-labeled objects and low-overlap objects) for group adjudication. The output of this workflow is a set of consensus-validated annotations suitable for supervised learning pipelines (14,16,17,23–25,36). Errors propagate even in large training datasets through to application, and thus Anatolution requires that each retained annotation reflects agreement among multiple trained observers and is cross-checked against computational detection for completeness.

Model predictions trained on consensus labels are subsequently validated against held-out consensus annotations, and systematic prediction errors are used to prioritize new annotation and adjudication—closing the loop between expert judgment and computational pattern recognition.

### 2.5 Segmentation pipeline

The segmentation pipeline produces computational “seeds”—candidate cell locations with centroid coordinates and contour boundaries—that serve as the objective backdrop against which human annotators work in the Anatolution interface. By providing an exhaustive catalogue of candidate objects, the seeds ensure that annotators evaluate a standardized set of structures, enabling systematic comparison across users and verification of annotation completeness. The centroid coordinates serve as the spatial frame of reference for matching annotations across users during consensus derivation.

The pipeline operates on individual 2D image tiles to generate new data. Specifically, the protocol proceeds through a sequence of local contrast enhancement, morphological smoothing, intensity thresholding, and local minima detection, with parameters adapted to the properties of each image. For Nissl-stained tissue, where neurons appear as dark, roughly elliptical structures against a lighter neuropil background, the pipeline operates on the value channel of the HSV color space, where cresyl violet absorption contrast is strongest. Contrast-limited adaptive histogram equalization (CLAHE; kernel size = 10, clip limit = 0.01, 100 bins) is applied to normalize local intensity distributions across images with variable staining quality and uneven illumination. The equalized image is inverted so that cell bodies are represented as high-intensity objects against a low-intensity background. Bilateral mean smoothing (disk radius = 50 pixels, intensity range parameters s0 = 100, s1 = 100) reduces within-object pixel variation while maintaining edge contrast at cell boundaries. Objects are retained by range thresholding at 50% of the image dynamic range (threshold = max − 0.50 × [max − min]), discarding low-intensity background while preserving faint but genuine cellular profiles. The thresholded image is inverted, and local minima—corresponding to the intensity centers of cell body regions—are detected with full connectivity. These minima are dilated using an elliptical structuring element (5 × 5 pixels, 5 iterations) to produce seed regions of sufficient spatial extent for contour extraction. A separate mask dilation (elliptical structuring element, 10 × 10 pixels, 1 iteration) provides a broader region for boundary delineation. Contours are extracted from the dilated seed regions using OpenCV’s contour detection with full hierarchy retrieval, and centroid coordinates are computed from image moments for each detected object. In immunohistochemical preparations like NeuN (9,11,40,41), the pipeline is merely adjusted to accommodate the alternative visual properties of the images under investigation to provide an adaptive specificity gate that helps accommodate variation in staining intensity across preparations. The pipeline’s morphological filtering stages smooth aggressively to suppress background noise and consolidate cell representations. This smoothing reduces fine morphological detail, including the crenelated soma boundaries characteristic of Nissl-stained neurons that reflect rough endoplasmic reticulum distribution. The detection stage therefore prioritizes reliable identification of cell locations over preservation of morphological detail—an appropriate tradeoff for a seed generation system whose purpose is to catalogue what structures exist and where they are located, while leaving morphological classification to human annotators. Fine boundary features are available in the raw imagery that annotators view during the labeling process and are captured separately in downstream feature extraction.

In support of three-dimensional annotation, detections from serial sections within a defined 3D image volume are linked across the Z-axis by spatial proximity in the XY-plane. For each contour detected in a given section, all contours in the immediately adjacent section whose centroid falls within ±60 pixels in both X and Y are assigned the same column identifier, and group membership is propagated iteratively through the depth of the volume. This proximity-based linkage produces columnar groupings of detections that approximate the through-plane extent of individual cells or cell clusters, enabling annotators to track objects across sections in the three-dimensional annotation interface. Bounding box coordinates for each column are computed as the extremes of constituent centroid positions, padded by 10% of the image dimensions (160 pixels in X, 120 pixels in Y for 1600 × 1200 pixel tiles), and expressed as percentages of image dimensions to maintain resolution independence across display contexts.

In parallel with seed generation, a separate preprocessing module produces standardized multiscale representations of each image tile for use in downstream supervised classification. This module operates independently of the seed generation pipeline and produces the engineered feature arrays against which consensus-validated annotations are evaluated in statistical learning workflows. Raw RGB images are converted to HSV color space, and three channels—hue, saturation, and value—are extracted as approximately uncorrelated feature dimensions capturing complementary aspects of staining intensity and chromaticity. Each channel is processed at three spatial scales using Gaussian filtering (sigma = 0, 20, and 50 pixels), producing nine feature arrays representing tissue structure from fine cellular detail to coarse regional variation. All nine channel-scale arrays are normalized using CLAHE (kernel size = 10, clip limit = 0.001, 125 bins) to standardize local contrast. A mean bilateral filter is applied within disk-shaped structuring elements scaled to each spatial frequency tier (radii of 10, 30, and 50 pixels). For the value channel, the bilateral filter’s intensity range parameters are inverted between Nissl preparations (s0 = 10, s1 = 100, preferentially smoothing bright background while preserving dark cell bodies) and NeuN preparations (s0 = 100, s1 = 10, preferentially smoothing dark background while preserving bright labeled profiles), ensuring that object boundaries are preserved regardless of stain polarity. Hue and saturation channels are filtered symmetrically (s0 = s1 = 25). A modal filter within local neighborhoods further regularizes intensity values while respecting region boundaries.

Scharr gradient operators—selected for their superior rotational invariance relative to Sobel operators—are applied to the smoothed arrays to produce edge gradient maps for all nine channel-scale combinations. Edge maps are normalized by CLAHE, then clipped at the 75th percentile of nonzero values and rescaled to the unit interval via min-max normalization, suppressing extreme gradient responses while preserving the relative structure of biologically meaningful boundaries. This base extraction stage produces 18 feature arrays per tile where nine represent smoothed intensity across three color channels at three spatial scales, and nine represent the corresponding normalized edge gradients, stored as compressed NumPy archives for downstream consumption.

Capturing the local spatial context on which cell-type classification depends, the three unblurred HSV channels (hue, saturation, and value at sigma = 0) serve as input for context computation. For each of these three channels, local mean and local variance are computed within disk-shaped kernels at three radii (3, 7, and 15 pixels, corresponding to kernel sizes of 7 × 7, 15 × 15, and 31 × 31 pixels)—spatial extents that approximate subcellular, cellular, and pericellular scales, respectively. This produces 18 context features per pixel (3 channels × 3 radii × 2 statistics). The context features encode how each pixel’s intensity and chromaticity relate to their immediate spatial surroundings—information that is critical for distinguishing cell types whose defining morphological features include not only intrinsic staining properties but also the contrast and texture of the surrounding neuropil. The full feature representation thus comprises 36 channels per pixel: 18 base channels encoding multiscale intensity and edge structure, and 18 context channels encoding local neighborhood statistics at biologically motivated spatial scales. All feature arrays are extracted at the native resolution of the input imagery (1600 × 1200 pixels for TIF; 960 × 720 pixels for JPG) and stored as compressed archives organized by container, ensuring that the feature representation is resolution-matched to the annotation coordinate system. Annotation coordinates, stored as percentages of image dimensions, are mapped to pixel coordinates at the native resolution of each image format during training data preparation, preserving spatial correspondence between labels and features regardless of the source format.

### 2.6 Downstream pipeline and results export

Annotation data are stored in a relational PostgreSQL database that preserves the relational structure of annotation vocabulary to image dimensionality. Data are stored to preserve consensus verification, including user identity, project context, image identifiers, marker type, and spatial coordinates recorded as percentages of image dimensions. For three-dimensional contour annotations, the database stores polygon boundaries as ordered coordinate sequences across Z-slices, enabling reconstruction of volumetric objects. In constructing volumetric morphometrics, attention must be paid to the spatial dimension represented by sequential images in a volume (i.e., the interval or distance between images), and the database preserves these relations for adjustment according to project demands. This relational structure ensures that per-user and per-project isolation is maintained, supporting the independent annotation and subsequent consensus computation that the workflow requires.

Data exports are generated via graphic user interface in common format. As common export patterns emerge from community use, user-facing interfaces for frequent operations may be added, but the underlying principle—that researchers retain full programmatic access to their data—will remain central to the platform’s design. Exported annotations are formatted to support direct ingestion by supervised learning pipelines. The GitHub repository includes preprocessing code to convert exported coordinate lists into image masks compatible with commonly used deep learning architectures, including nnU-Net (36) and Mask R-CNN (42,43). Annotation coordinates are stored as resolution-independent percentages of image dimensions, and thus masks can be generated at any target resolution appropriate to the training regime. The complete pipeline from consensus-validated annotations to model training data is documented in the repository, with the goal of minimizing the technical friction between annotation and computation for the community moving forward.

### 2.7 Validation corpus

The validation analyses reported here draw on a corpus of polygon annotations collected via the Anatolution platform from primate histological preparations. All specimens were drawn from non-human primate specimens in the Kaas collection as part of DJM’s dissertation (44). A total of 48 raters contributed annotations to the platform. Of these, 39 had sufficient data for computation of a composite Reliability Index (RI = 0.4 × Agreement + 0.3 × Stability + 0.3 × Coverage), where Agreement is the median majority-vote Dice across container–marker combinations, Stability is the inverse of cross-container standard deviation of majority-vote Dice (normalized), and Coverage is the fraction of total containers annotated (out of 18 maximum). Among these 39 raters, 28 annotated across multiple containers and 11 contributed to a single container.

Tissue sections were digitized as full-resolution tiles (1600 × 1200 pixels for TIF; 960 × 720 pixels for JPEG) from archived specimen collections (45). Statistics from as many as five staining categories were summarized in this report spanning approximately 10 years of work (9) spanning a collection that took 5 decades or work (46) to generate: Nissl (n = 433 tiles), Nissl-NeuN double-stain (NsNn; n = 232), medial geniculate nucleus with biotinylated dextran amine tract tracing (MGN/BDA; n = 163), Myelin (n = 151), and NeuN immunohistochemistry (n = 45), totaling 1,024 tiles across 81 containers. A container represents a single histological probe—a column-shaped tissue region extracted from one section of one brain for one staining preparation—and serves as the fundamental organizational unit for both annotation and analysis. Polygon annotations were collected from 48 raters, of whom 23 contributed across multiple containers and 25 annotated within a single container. Target cell-type markers included Neuron, Astrocyte, Oligodendrocyte, Microglia, Endothelial, and Artefact, with an additional Unknown marker available for cases of low classification confidence. The master annotation dataset contained 603,297 polygon annotation rows spanning the full corpus. Containers were classified into three quality tiers based on rater coverage, following the principle that consensus reliability increases with annotator count. Gold-tier containers (n = 5; 46 tiles; ≥7 independent raters each) provided the highest-confidence consensus estimates and served as the primary validation dataset. Silver-tier containers (n = 14; 123 tiles; ≥3 raters each) provided secondary validation and supplementary training data. Bronze-tier containers (n = 9; 1–2 raters) were used for quality characterization but were not eligible for consensus-based model training. This tiered structure reflects a deliberate design choice: rather than treating all annotations as equivalent, the platform stratifies data by the number of independent observations available, enabling users to select training data appropriate to the reliability level their application demands.

### 2.8 Analytical methods

We used conventional rater reliability metrics to assess data quality (38,47). Specifically, we evaluated inter-rater reliability, consensus quality, seed detection performance, rater quality, and computed an automated baseline. All analyses were implemented in Python using NumPy, SciPy, and scikit-learn, with code available in the project repository.

Inter-rater boundary agreement was quantified using the Dice coefficient (also known as the Sorensen-Dice coefficient), computed for each rater-pair combination within each container-marker-tile combination (24,36). For a given tile and marker, each rater’s polygon annotations were rasterized into a binary mask at the native image resolution. The Dice coefficient between raters i and j was computed as 2|A_i_ intersect A_j_| / (|A_i_| + |A_j_|), where |A| denotes the number of foreground pixels. For containers with n raters, all C_(n,2)_ pairwise comparisons were computed, yielding a distribution of Dice values per container-marker combination. Median, interquartile range (IQR), and mean Dice were computed across all tile-marker-pair comparisons within each stratum (by marker type, by stain type, and overall).

Two ceiling estimates were computed for each container-marker combination to bound the achievable agreement between an individual rater and the consensus target (48). The majority-vote (MV) ceiling was computed by generating a pixel-wise majority-vote consensus mask from all n raters, then computing the Dice coefficient between each rater’s mask and this consensus. Because each rater contributes to the consensus target, the MV ceiling is positively biased and represents an upper bound. The leave-one-out (LOO) ceiling was computed by excluding each rater in turn from the consensus computation, generating a majority-vote mask from the remaining n-1 raters, and computing the Dice coefficient between the excluded rater’s mask and this held-out consensus. The LOO ceiling provides a stricter, approximately unbiased estimate of how well an individual rater agrees with the consensus of others.

We subsampled top-tier quality annotations to show the relationship between consensus and individual accuracy in labeling cells. Specifically, up to 200 random subsets of k raters were drawn without replacement for each value of k (from 1 to the maximum number of raters in the container). For each subset, a pixel-wise majority-vote consensus mask was computed and compared to the full-rater consensus via Dice coefficient. Median Dice and interquartile range across tiles were computed for each value of k, producing a convergence curve to quantify the marginal improvement in consensus stability per additional rater. To characterize individual rater quality across the corpus (47), a composite Reliability Index (RI) was computed for each rater with sufficient annotation data. The RI is defined as a weighted sum of three components: RI = 0.4 × Agreement + 0.3 × Stability + 0.3 × Coverage, where Agreement is the median majority-vote Dice across all container-marker combinations annotated by the rater, Stability is the inverse of the cross-container standard deviation of majority-vote Dice (min-max normalized across raters), and Coverage is the fraction of total training-eligible containers (Gold and Silver tier; n = 18 maximum) annotated by the rater. The weighting reflects the priority assigned to boundary agreement as the primary quality criterion, with stability and breadth of engagement as secondary indicators.

During weighted consensus generation, per-rater quality weights were derived from leave-one-out (LOO) Dice distributions. For each rater-marker pair, the median LOO Dice across tiles was computed, and weights were assigned proportional to this median. Weights were bounded to the interval [0.5, 1.5] and normalized within each consensus set so that they sum to the number of raters. This bounding ensures that high-agreement raters exert proportionally greater influence on the final consensus mask while maintaining a floor that prevents any rater’s contribution from being discounted entirely.

The segmentation pipeline was evaluated in two complementary modes: a conservative L2 mode optimized for precision (fewer seeds, higher specificity) and a high-recall M5 mode optimized for completeness (more seeds, lower specificity). Performance was assessed using both pixel-overlap metrics and spatial anchoring metrics (48). Pixel-overlap metrics included Dice coefficient, precision, and recall between the binary seed mask and the consensus annotation mask. Spatial anchoring metrics included the seed inside rate (fraction of detected seeds whose centroid falls within any annotated polygon boundary) and the polygon coverage rate (fraction of annotated polygons containing at least one seed centroid). A composite objective mismatch score was computed per tile as a weighted combination of (1 - seed inside rate), (1 - polygon coverage rate), and overdensity (the ratio of seed count to polygon count, clipped at 1.0), providing a summary measure of annotation-to-seed alignment.

A pixel-level Random Forest classifier (Context-RF) was trained using the 36-channel multiscale feature representation described in Section 2.5 as input and the majority-vote consensus mask as the training target. The classifier was configured with 200 trees and a maximum depth of 20. Performance was evaluated via leave-one-tile-out cross-validation within each container: for each tile, the classifier was trained on all other tiles in the container and predicted the held-out tile. Dice coefficients between the predicted mask and the majority-vote consensus mask were computed for each held-out tile and compared to the MV and LOO ceiling estimates to establish the relative performance of the automated baseline against the human annotation ceiling (47,48). To evaluate whether consensus-derived annotations support end-to-end supervised learning, we export data in preparation for common biomedical image pipelines like the nnU-Net framework (36). Briefly, images are stored as channels aligned with their labels as 2D tiles for ingestion and training.

## Results

Forty-eight independent raters produced 603,297 polygon annotations across 1,024 histological tiles using the Anatolution platform (Figure 1). Annotations were derived primarily from two staining categories — Nissl (n = 433 tiles) and NsNn (n = 232 tiles) — which together constitute the primary analytical corpus because both preparations label all cell types simultaneously, enabling cross-class annotation comparisons within each tile. Within the Nissl and NsNn corpus, we report results for six cell-type markers: Neuron, Astrocyte, Oligodendrocyte, Microglia, Endothelial, and Artefact (Table 1). Of the 81 containers in the corpus, 19 met criteria for inclusion in the training-eligible Gold and Silver tiers: 5 Gold-tier containers (46 tiles, ≥7 independent raters each) and 14 Silver-tier containers (123 tiles, ≥3 raters each). The remaining containers were classified as Bronze (1–2 raters) and used for quality characterization but not for consensus training data.

**Figure 1.**
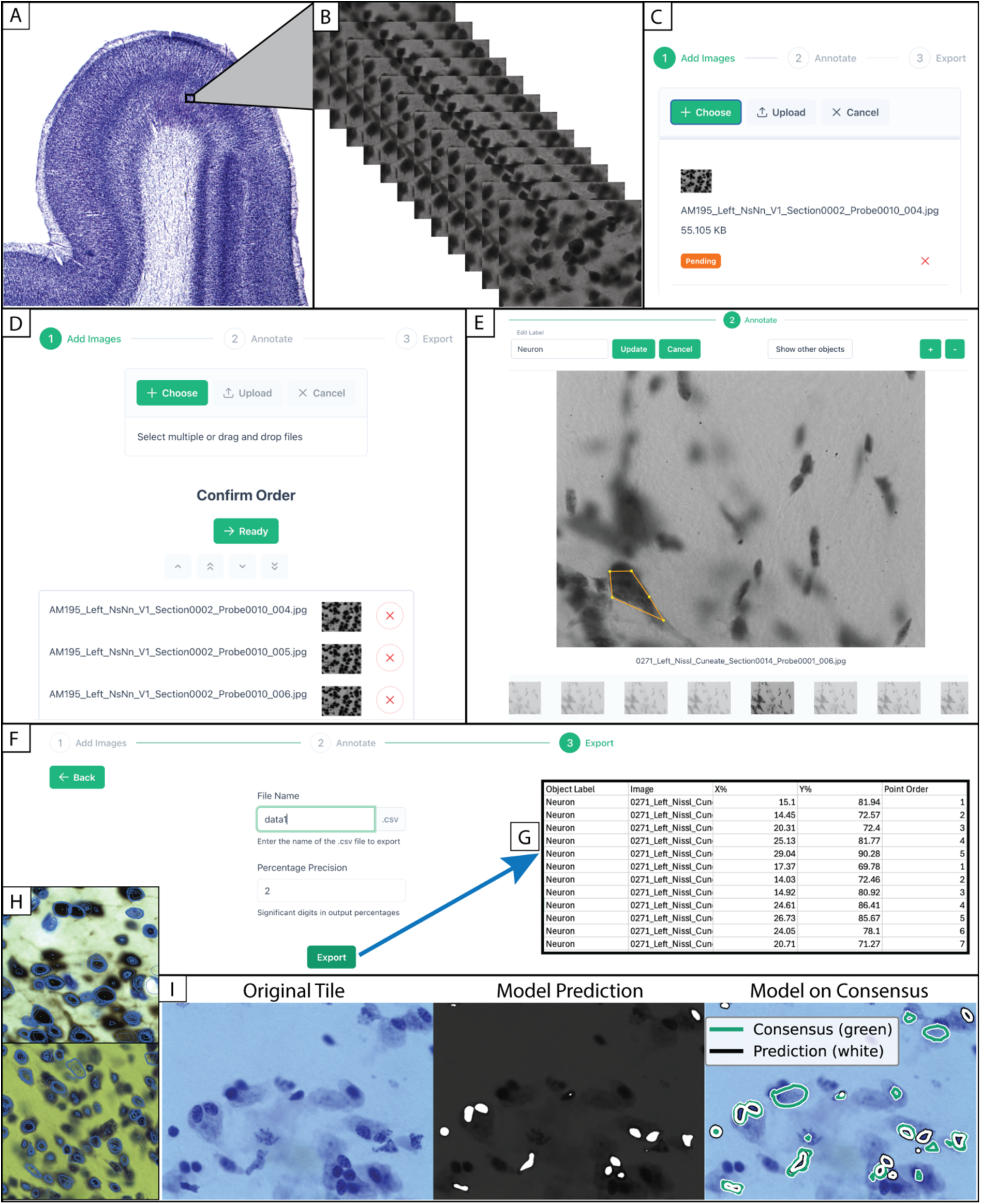
Overview of the Anatolution Platform. Overview of the Anatolution platform and GUI, illustrating the end-to-end pipeline from histological image acquisition through multi-rater consensus annotation to downstream computational model evaluation. The figure is organized in three tiers: image ingestion and three-dimensional tissue representation (A–B), the browser-based annotation workflow including image upload, polygon delineation, and structured data export (C–G), and preliminary model training with consensus-based validation (H–I). (A) Light microscopy image of a Nissl-stained coronal section from a non-human primate brain (*Papio anubis*). The boxed region indicates a single probe location selected for annotation. (B) Serial focal planes through the selected tissue region, illustrating the volumetric probe structure used for cellular annotations. Each probe comprises 13–17 focal planes acquired at approximately 1 µm axial spacing (~15 µm total depth), which annotators navigate along the Z-axis to delineate cellular structures across depth (see Methods §2.3). (C) Image upload interface, showing step 1 of the 3-step annotation workflow. A single NsNn tile is staged for upload using a file naming convention that encodes the hierarchical image organization used by the platform for directory structure and project management. (D) Image ordering interface, in which uploaded tiles are arranged into the correct Z-dimension sequence prior to annotation. Users confirm the serial order of focal planes within each probe using the sorting controls shown to define any Z-stack through which annotators navigate during polygon delineation. (E) Annotation interface showing step 2 of the workflow where a Nissl-stained tissue tile is displayed with a completed polygon annotation of a neuronal soma (yellow boundary). The label editor (top) shows the current marker assignment (“Neuron”), and the Z-stack navigation strip (bottom) displays thumbnail previews of serial focal planes, enabling annotators to trace object boundaries across images in a volume. Configurable marker vocabularies (e.g. Neuron) are defined in the GUI, and individual polygons are stored and retrieved independently to preserving independence for consensus derivation. (F) Data export interface showing step 3 of the workflow to get data (G). Completed annotations are exported as CSV files containing object labels and polygon coordinates for reconstruction and ingestion with conventional bioimage pipelines. (H) Representative Nissl-stained tissue tile(s) at high magnification, showing individual drawings (blue lines) of neuronal and glial cell bodies against computer vision seeds (yellow dots). (I) Preliminary downstream model validation on a top-tier container (Nissl stain of cuneate nucleus with 12 independent raters). Original color tile on the left followed by overlay of model in the center and the overlay of model (white) on consensus labels (green) at right. The two features that distinguish Anatolution from existing toolkits are represented across this figure from project initiation (A, B) to the structured multi-rater annotation workflow (C–G), in which independent annotators label identical image sets against a shared computational backdrop to produce consensus-validated ground truth, and the integration of that consensus output with downstream supervised learning (I), closing the loop between expert morphological judgment and computational pattern recognition.

**Table 1.** Consensus of users across object marker types. Pairwise boundary Dice coefficients across 17,982 tile–marker–pair comparisons from 31 containers with two or more independent raters. Dice coefficients were computed for all rater-pair combinations within each container–marker–tile combination. Oligodendrocyte markers showed the highest median agreement, consistent with the compact, darkly staining morphology that produces well-defined boundaries in Nissl preparations. Neuron markers, which constituted the largest share of comparisons, showed moderate agreement consistent with the recognized difficulty of delineating neuronal somata in densely packed cortical regions. Artefact markers exhibited the widest variability (IQR = 0.804), reflecting the heterogeneity of structures assigned to this catch-all quality-control category. NsNn-stained tiles showed higher median Dice than Nissl-stained tiles, likely reflecting sharper chromatic contrast and more compact cell profiles in the double-stained preparations.

| *Marker* | *n_Pairs* | *Median_Dice* | *Q1* | *Q3* | *IQR* | *Mean_Dice* |
| --- | --- | --- | --- | --- | --- | --- |
| *Oligodendrocyte* | 3284 | 0.498 | 0.189 | 0.74 | 0.551 | 0.464 |
| *Endothelial* | 1941 | 0.476 | 0.139 | 0.752 | 0.613 | 0.451 |
| *Microglia* | 1439 | 0.441 | 0.154 | 0.707 | 0.553 | 0.442 |
| *Astrocyte* | 4054 | 0.427 | 0.144 | 0.664 | 0.52 | 0.411 |
| *Neuron* | 6139 | 0.425 | 0.146 | 0.683 | 0.537 | 0.415 |
| *Artefact* | 897 | 0.328 | 0 | 0.804 | 0.804 | 0.391 |
| *Overall* | 17982 | 0.444 | 0.139 | 0.707 | 0.568 | 0.428 |

We computed consensus Dice coefficients against the full-rater reference for k-rater subsamples across Gold-tier containers to quantify the relationship between annotator count and consensus quality (Figure 2). Consensus quality increased monotonically with annotator count from a median consensus Dice at approximately 0.30 at k = 1 to 0.63 at k = 3, 0.79 at k = 7, and approached 1.0 as k approached the full rater pool. The rate of improvement diminished beyond approximately seven annotators, with the consensus Dice curve approaching an asymptotic plateau.

**Figure 2.**
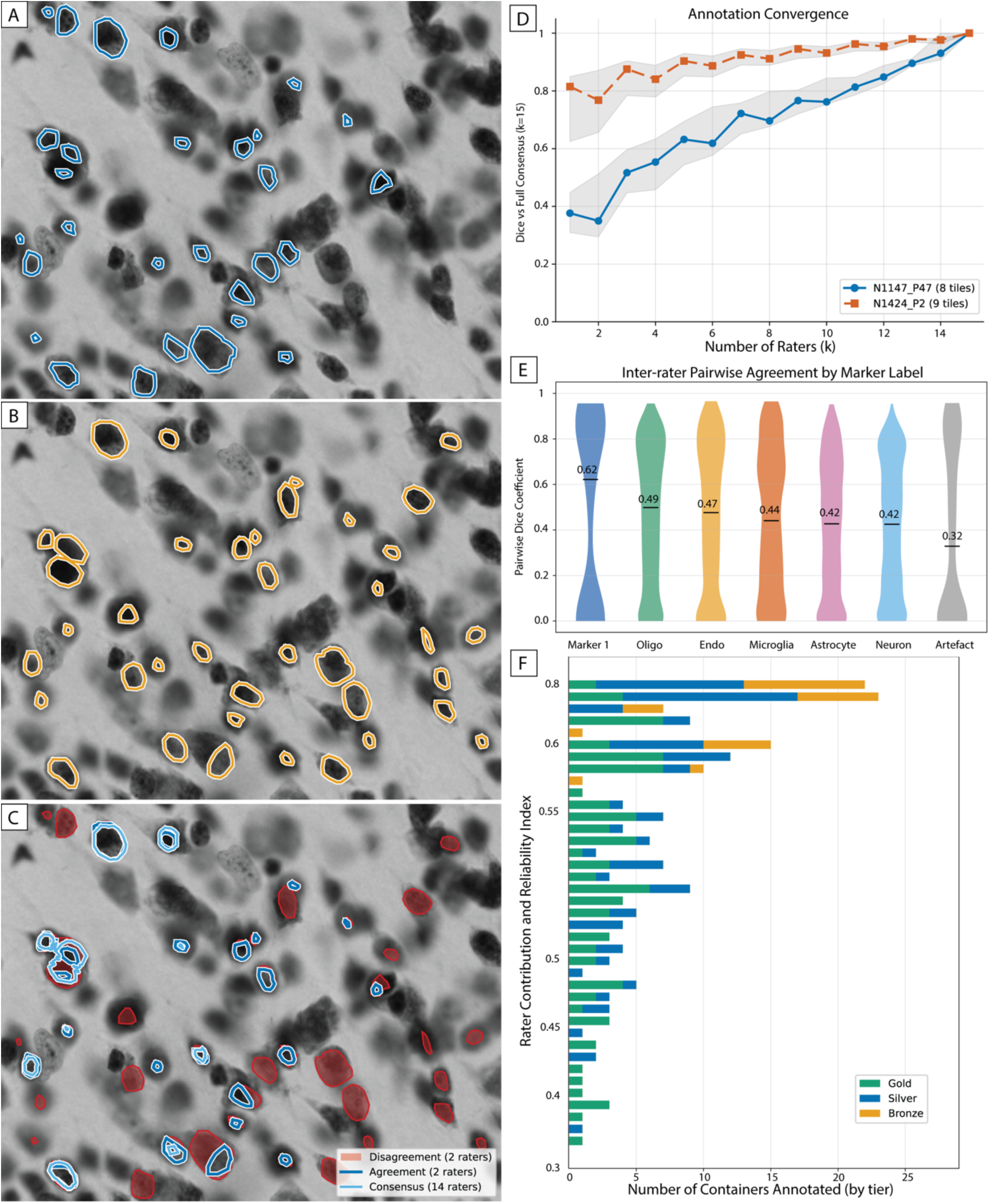
Evaluation of Consensus Behavior. This figure presents the multi-rater consensus process and its quantitative validation across six panels. Panels A–C illustrate the consensus workflow on a single top-tier tile, showing how independent annotations from multiple raters are combined into a consensus boundary and how agreement and disagreement are visualized. Panels D–F summarize corpus-wide metrics for annotation convergence, inter-rater boundary agreement by marker type, and per-rater reliability and contribution. Panel (A) shows the independent polygon annotations from a first and then (B) a second user followed by (C) showing agreement and disagreement between raters overlaid by the consensus label. Agreement regions (intersection of user polygons) are shown as dark blue contours while disagreement regions (symmetric difference) are shown as red (contour and fill), and the majority-vote consensus boundary derived from all 14 raters is shown as light blue contours. (D) Annotation convergence curve showing the Dice coefficient between subsampled consensus (k raters, where k = 1–15) and full consensus (k = 15) for the Neuron marker in two top-tier containers. For each value of k, up to 200 random rater subsets were drawn and majority-vote consensus was computed at the pixel level. Lines show the median Dice across tiles and shaded bands show the interquartile range. At k = 3 raters, median Dice reaches 0.629; at k = 7 (the Gold-tier threshold), median Dice reaches 0.792; at k = 15, Dice = 1.0 by definition. The curve demonstrates the need for ~7 annotators and suggests diminishing returns ≥7 annotators for ceiling-stable consensus estimates. (E) Distribution of pairwise Dice coefficients by marker type across 17,982 tile–marker–pair comparisons from 31 containers with ≥2 raters. Each violin shows the full distribution; horizontal bars indicate median values. Marker 1 (median Dice = 0.62; n = 228 pairs) showed the highest agreement, followed by Oligodendrocyte (0.49; n = 3,284), Endothelial (0.47; n = 1,941), Microglia (0.44; n = 1,439), Astrocyte (0.42; n = 4,054), Neuron (0.42; n = 6,139), and Artefact (0.32; n = 897). (F) User Reliability Index (RI) and contribution to containers at each quality tier for 39 raters with sufficient data for RI computation. Each horizontal bar represents one rater, ordered by RI (y-axis). Bar segments are colored by the number of containers annotated at each tier: Gold (green; ≥7 raters per container), Silver (blue; ≥3 raters), and Bronze (orange; 1–2 raters). RI is a composite score defined as 0.4 × Agreement + 0.3 × Stability + 0.3 × Coverage, where Agreement is the median majority-vote Dice across container–marker combinations, Stability is the inverse of the cross-container standard deviation of majority-vote Dice (normalized), and Coverage is the fraction of total training-eligible containers annotated. Mean RI across raters was 0.514 (SD = 0.109; range: 0.324–0.831; median: 0.503). The highest-performing raters (RI > 0.6) contributed across multiple tiers and annotated the largest number of containers, demonstrating both sustained engagement and consistent quality. Raters with lower RI values (below 0.40) tended to contribute to fewer containers and showed higher variability in boundary placement. Panels A–C demonstrate the central methodological contribution of Anatolution to enable independent annotations from trained raters, collected without mutual visibility, to converge on consistent object boundaries when aggregated through consensus. Panel D quantifies this convergence, showing that consensus stability increases monotonically with rater count and approaches an asymptote near k = 7. Panels E and F contextualize these findings across the full corpus, revealing that agreement varies systematically by marker type and object difficulty.

**Figure 3.**
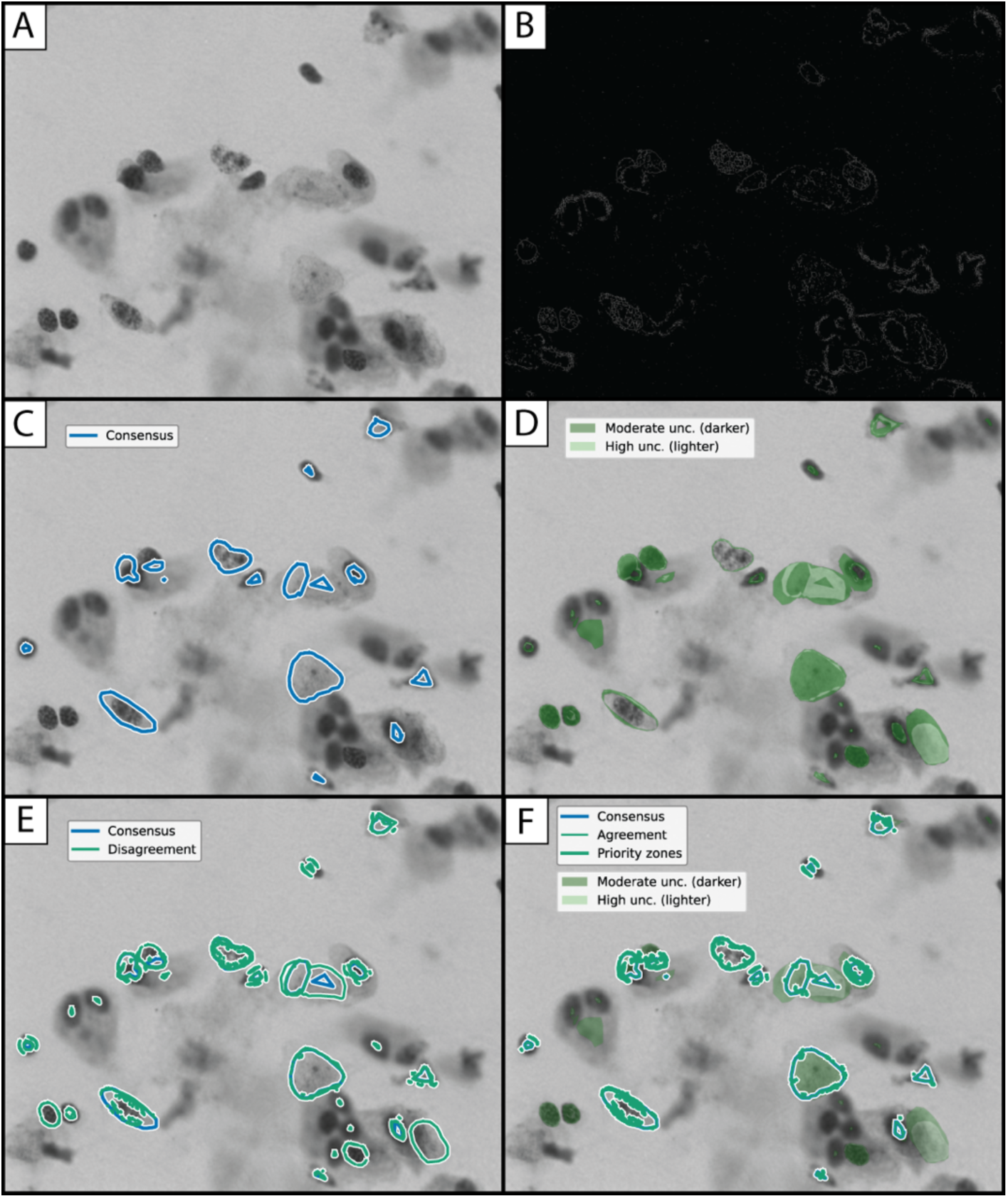
Augmented decision support. Six-panel demonstration of the decision-support overlays that augment human consensus labels with image-derived structural information. All panels display the same tissue in grayscale as background (A), or edges calculated on this image (B), where white lines on the black background correspond to sharp intensity transitions in the tissue — typically cell-body edges, vessel boundaries, or staining artifacts. Stained cell bodies appear as dark structures against a lighter neuropil background, and (C) shows consensus labels alongside (D) depicting uncertainty among users (uncertainty increases from dark to light green shading). Panel (E) shows consensus in blue against red disagreement alongside the composite decision-support overlay (F) showing the relationship among measures of consensus and uncertainty. Priority review zones were identified within the 30-pixel consensus boundary band by computing z-scores for both uncertainty and edge magnitude; pixels where z(uncertainty) + z(edge) > 2.0 were flagged. These zones mark the highest-priority regions for manual boundary review — locations where both rater disagreement and tissue structure jointly indicate boundary ambiguity. All priority review flags fall within the consensus boundary band.

We calculated pairwise Dice coefficients to assess whether the platform’s per-container metrics discriminate annotation quality across morphological conditions. We found that all C_(n,2)_ rater-pair combinations across 31 containers with two or more raters yielded 17,982 tile–marker–pair comparisons. Per-container reliability metrics revealed systematic variation in achievable agreement that tracked both marker type and annotation maturity (Table 2). Containers targeting well-defined structures with adequate rater coverage achieved agreement consistent with the convergence curve’s asymptotic plateau: N1424_P2 (Oligodendrocyte, 15 raters, Gold) reached a median pairwise Dice of 0.735, and N1424_P5 (Artefact, 6 raters, Silver) reached 0.923 (Table 2). Even below the Gold-tier threshold, IB290 (Neuron, 4 raters, Silver) achieved a leave-one-out ceiling Dice of 0.666 ± 0.024. At the other extreme, N1424_P3 (Neuron, 11 raters, Gold) showed a median pairwise Dice of 0.088 despite one of the largest rater pools in the corpus (Table 2).

**Table 2.** Consensus convergence and reliability. Per-container reliability profiles for the four extreme cases in the annotation corpus, illustrating how the platform’s per-container metrics discriminate annotation quality across morphological conditions and rater pools. Pairwise median Dice was computed across all rater-pair combinations within each container–marker combination. The LOO ceiling Dice was computed by excluding each rater in turn and measuring agreement against the remaining consensus (see Methods §2.8). Convergence status was assigned based on position along the empirical convergence curve (Figure 2D): “at asymptote” denotes containers whose agreement falls at or near the plateau observed beyond 7 raters; “approaching threshold” denotes containers with fewer than 7 raters whose LOO ceilings indicate convergent annotation; “boundary ambiguity detected” denotes containers where low agreement despite adequate rater coverage signals genuine morphological uncertainty rather than rater inconsistency. This classification reflects the platform’s design principle that per-container reliability metrics should inform downstream data stratification rather than serve solely as summary quality indicators.

| Marker | Raters | Tier | Pairwise<br>Median<br>Dice | LOO<br>Ceiling<br>Dice | Convergence<br>Status | Interpretation |
| --- | --- | --- | --- | --- | --- | --- |
| Oligo | 15 | Gold | 0.735 |  | At asymptote | Compact, darkly staining morphology with maximum rater pool; boundary agreement approaches clinical histopathology benchmarks. |
| Artefact | 6 | Silver | 0.923 |  | At asymptote | Highest pairwise agreement in the corpus on a morphologically unambiguous category. |
| Neuron | 4 | Silver |  | 0.666<br>±<br>0.024 | Approaching threshold | Strong LOO ceiling on a difficult cell type with a sub-Gold rater pool. |
| Neuron | 11 | Gold | 0.088 |  | Boundary ambiguity detected | Dense cortical neuropil where neuronal somata overlap and boundary placement is genuinely ambiguous. |

We estimated performance ceilings for each container–marker combination using two complementary approaches. The majority-vote (MV) ceiling — an upper bound in which each rater contributes to the consensus target — ranged from 0.227 to 0.790 (median Nissl = 0.566, median NsNn = 0.590). The leave-one-out (LOO) ceiling — a stricter lower bound that excludes the evaluated rater from the consensus — ranged from 0.000 to 0.769 (median Nissl = 0.319, median NsNn = 0.306). For the Neuron marker — the most morphologically challenging cell type in the corpus — the three highest-performing containers by LOO ceiling demonstrated that neuronal boundaries are tractable when annotators and tissue quality align: LOO = 0.677 ± 0.110 (15 raters), LOO = 0.666 ± 0.024 (4 raters), and LOO = 0.621 ± 0.218 (12 raters).

We trained a pixel-level Random Forest classifier (Context-RF on 36 HSV-derived features, max depth = 20 for 200 trees) using leave-one-tile-out cross-validation within each container to establish an automated baseline for comparison against human labels. The Context-RF achieved 50–56% of the MV ceiling on average, indicating that a simple classifier operating on local color and texture features captures roughly half the consensus signal.

We measured the correspondence between algorithmically detected seeds and human-annotated polygon boundaries on 68 tiles (45 Nissl, 23 NsNn) to evaluate the segmentation pipeline’s role as a spatial anchoring system. In conservative L2 detection mode (median 73 seeds per tile for Nissl; 58 for NsNn), 88.1% of annotated polygons contained at least one algorithmically detected seed, and 37.1% of L2 seeds fell within annotated polygon boundaries (Table 3). To assess whether the pipeline can provide an exhaustive completeness check, we evaluated the high-recall M5 detection mode, which produces an order of magnitude more seeds per tile than L2 (median 1,965 for Nissl; 553 for NsNn). M5 achieved near-complete polygon coverage for Nissl tissue (100% median) at the cost of a lower seed-inside rate (9.6%), consistent with its design as a sweep that prioritizes detecting every cellular structure over suppressing false positives. A composite objective mismatch score provided a single per-tile quality indicator for automated flagging of tiles requiring re-examination and showed Nissl tiles at a median mismatch of 0.282, while NsNn tiles showed a lower median mismatch of 0.193, providing a per-tile indicator that complements rater reliability.

**Table 3.** Comparison of features to literature. Qualitative feature comparison of Anatolution with established bioimage analysis platforms. Features are grouped by functional category: annotation workflow, consensus and quality control, segmentation capability, data management, and deployment. This comparison is intended to illustrate the complementary roles of these tools rather than to suggest competition: CellProfiler, Ilastik, Cellpose, and QuPath address algorithmic challenges in segmentation and classification, whereas Anatolution addresses the upstream constraint of producing consensus-validated training data for these and similar tools. Feature availability was assessed from published documentation and the papers cited (27–32) as of 2025 in case of changes pending. The central distinction illustrated by this comparison is architectural rather than algorithmic. CellProfiler, Ilastik, Cellpose, and QuPath each provide powerful capabilities for image analysis, segmentation, and classification — capabilities that depend on the availability of training and evaluation data adequate to the target domain. Anatolution occupies a distinct position within this larger system that is upstream of some tools and largely complementary to the existing active statistical learning ecology. Specifically, this report provides the infrastructure to produce consensus-validated labels through a structured multi-annotator workflow with integrated reliability assessment that is generally machine-readable. The platform’s outputs are designed to be consumed by any downstream framework, especially those listed in this report. In this sense, Anatolution does not compete with existing tools but addresses the prerequisite that limits their effective deployment in domains where single-annotator labels are insufficient, as the field has discovered (14).

| *Feature* | *Anatolution* | *CellProfiler* | *Ilastik* | *Cellpose* | *QuPath* |
| --- | --- | --- | --- | --- | --- |
| *Multi-annotator support* | Yes | No | No | No | No |
| *Per-user annotation isolation* | Yes | No | No | No | No |
| *Configurable marker vocabulary* | Yes | No | Yes | No | Yes |
| *Polygon boundary annotation* | Yes | No | Limited | Limited | Yes |
| *3D image volume annotation* | Yes | No | Yes | Limited | Limited |
| *Resolution invariance* | Yes | Limited | No | Limited | No |
| *Consensus label derivation* | Yes | No | No | No | No |
| *Rater reliability and weighting* | Yes | No | No | No | No |
| *Tiering by rater quality* | Yes | No | No | No | No |
| *Convergence analysis* | Yes | No | No | No | No |
| *Built-in pipeline* | Pixel/object | Modular | Pixel/object | Deep learning | Cell detection |
| *Pre-trained deep learning models* | No | Plugins | No | Yes | Plugins |
| *Interactive statistical learning* | Programmatic | No | Yes | No | No |
| *Whole-slide image support* | Yes | Yes | Yes | Limited | Yes |
| *Relational database* | Yes | No | No | No | No |
| *Programmatic data export* | Yes | Yes | Yes | Yes | Yes |
| *Per-project data isolation* | Yes | No | Yes | No | Yes |
| *Web-based (browser access)* | Yes | Limited | No | No | Limited |
| *Primary purpose* | Consensus | Modular | Interactive | Cell detection | Digital pathology |
| *Pipeline position* | Label production | Measurement | Classification | Detection | Analysis |
| *Primary output* | Consensus Annotations | Quantitative measures | Pixel/object classifier | Cell masks | Annotated measures |
| *Target user* | Programmer | Bioimage analyst | Bioimage analyst | Bioimage analyst | Pathology, student |
| *Open source* | Yes | Yes | Yes | Yes | Yes |
| *Primary references* | Current | (27,29,30) | (31) | (32,51) | (52) |

The Reliability Index (RI) revealed a continuous distribution of annotation quality across the 48-rater pool. Mean RI was 0.514 (range: 0.324–0.831), with RI values distributed continuously rather than clustering into discrete skill categories. The top-performing raters demonstrated both high agreement with consensus and broad engagement across the corpus (Figure 2), while low RI values (below 0.40) were associated with fewer containers and showed higher variability in boundary placement.

## Discussion

Our results demonstrate that the Anatolution platform produces consensus annotations whose quality is measurable, improvable with additional annotators, and tractable for downstream model training. Forty-eight independent raters contributed 603,297 polygon annotations across 1,024 histological tiles spanning five stain types, and inter-rater boundary agreement—quantified by pairwise Dice coefficients—reached a median of 0.444 across the corpus and exceeded 0.7 in the highest-quality Gold-tier containers with seven or more annotators. Consensus quality increased monotonically with annotator count, reaching a median Dice of 0.79 against the full-rater reference at seven raters, with leave-one-out ceiling Dice values of 0.621–0.769 for Neuron markers in Gold containers demonstrating that consensus estimates converge toward stable ground truth as the annotator pool grows. The segmentation pipeline detected candidate cell bodies with 88% polygon coverage against consensus-validated annotations, a pixel-level Random Forest classifier achieved 50–56% of the majority-vote ceiling, and the Reliability Index (RI) revealed a continuous distribution of rater quality (mean RI = 0.514, range 0.324–0.831) that the platform’s weighted consensus mechanism is designed to exploit. These findings indicate that the platform performs as designed and that structured consensus annotation yields more consistent ground truth than any single annotator’s labels alone, not because individual annotators are unreliable, but because even modest disagreement among skilled observers compounds when labels are applied at scale.

The improvement in annotation quality with multiple annotators reflects a fundamental principle in measurement science. Specifically, that expert judgment contains both signal—accurate biological knowledge applied to morphological identification—and noise—individual bias, attention fluctuations, and criterion drift—and averaging across independent observers reduces the latter while preserving the former (3,8,35). This principle is well established in the stereological literature, where the single largest source of variance in morphometric datasets is often the investigator herself (3,9,14,23), and where intraclass correlation coefficients have long served as both quality control measures and indicators of dataset reliability (35,38,47). Anatolution operationalizes this principle for the specific context of supervised learning, where the consequences of annotation noise are not merely imprecise estimates but systematically degraded model performance across millions of applications. The empirical convergence curve—median consensus Dice rising from approximately 0.30 at one rater to 0.63 at three and 0.79 at seven—provides a quantitative basis for this design and supports the Gold-tier threshold of seven or more annotators as a practical criterion for ceiling-stable consensus.

The computational segmentation seeds are critical to realizing this measurement principle in practice. Providing an objective set of candidate objects with defined centroids and boundary contours in the seeds, we establish a shared reference frame that ensures annotators are evaluating the same structures, making inter-annotator comparison meaningful rather than confounded by differences in what was detected. Without this shared reference, freehand annotations may not refer to the same objects at all, and apparent disagreement between annotators may reflect differences in detection or completeness rather than differences in classification decisions. The 88% polygon coverage achieved by the conservative L2 seed detection mode confirms that the pipeline captures a majority of structures that trained annotators identify, fulfilling its intended role as a spatial anchoring system. The complementary metric—that only 37% of detected seeds fall within annotated polygons—is consistent with the known incompleteness of manual annotation relative to the total population of visible cell bodies and underscores the value of algorithmic detection as a completeness check that neither source alone could provide. The seeds thus serve a specific and limited function in that they standardize *what* is being evaluated without constraining *how* annotators classify it and thus preserve the independence of judgment on which consensus depends.

The overall median pairwise Dice of 0.444 warrants careful interpretation. This value aggregates comparisons across 31 containers spanning five stain types, six cell-type markers, and rater pools ranging from 2 to 15 annotators—a deliberately heterogeneous corpus that includes containers at every stage of the annotation lifecycle, from early-stage Bronze-tier collections to mature Gold-tier datasets. The variation in agreement across this corpus is itself informative: Oligodendrocyte markers showed the highest median agreement (0.498), consistent with the compact, darkly staining morphology that produces well-defined boundaries in Nissl preparations, while Neuron markers showed moderate agreement (0.425), reflecting the recognized difficulty of delineating neuronal somata in densely packed cortical neuropil where cell boundaries are often ambiguous even to experienced observers (3,9,38,47). The wide variability of Artefact markers (IQR = 0.804) reflects the heterogeneity inherent to a catch-all quality-control category that encompasses incomplete profiles, out-of-plane objects, and non-cellular structures—precisely the kind of boundary case that the ‘unknown’ label is designed to resolve. Briefly, this variation reinforces our approach to leverage complementary human and computational approaches to measure more systematically.

Stain-type differences in agreement are similarly interpretable given known challenges in neuroanatomy. NsNn double-stained preparations showed higher median Dice (0.490) than Nissl-only preparations (0.417) because the chromatic contrast provided by the NeuN immunolabel sharpens soma boundaries and produces more compact cellular profiles that are easier to delineate consistently. This observation suggests that staining quality is not merely a technical variable but a direct determinant of achievable inter-rater agreement, a relationship that the platform’s per-container reliability metrics can capture and that should inform the design of future annotation protocols. The highest container-level agreements—0.923 for Artefact markers in a well-characterized container and 0.735 for Oligodendrocyte in a 15-rater Gold container—demonstrate that the consensus workflow can produce boundary agreement approaching the levels typically reported for expert-expert comparisons in clinical histopathology (38,47), provided that sufficient annotators contribute and that the morphological targets are well defined (8,49).

The contrast between the highest- and lowest-performing containers illuminates a key property of the consensus approach. N1424_P2 (Oligodendrocyte, 15 raters, Gold) achieved a median pairwise Dice of 0.735, while N1424_P3 (Neuron, 11 raters, Gold) showed a median of 0.088 despite one of the largest rater pools in the corpus. This disparity—between two Gold-tier containers with comparably large rater pools—demonstrates that annotator count alone does not determine annotation quality but that morphological difficulty is a co-determinant (14,16,49,50). In densely packed cortical neuropil, neuronal somata overlap and boundary placement becomes morphologically indeterminate, producing genuinely low agreement that reflects biological ambiguity (9) rather than rater inconsistency. That the platform’s per-container reliability metrics detect this distinction—rather than treating all containers as equivalent—is a design feature, not a limitation. In particular, this enables downstream users to stratify training data by achievable quality rather than by annotator count alone. Critically, even for this morphologically challenging cell type, other containers demonstrated that neuronal boundaries are tractable when annotators and tissue quality align, with leave-one-out ceiling Dice values reaching 0.666 ± 0.024 in a four-rater Silver container.

The continuous distribution of rater quality revealed by the Reliability Index (mean RI = 0.514, range 0.324–0.831) confirms that annotation skill is not a binary attribute but a graded capacity that varies across individuals and develops over time as is conventionally understood. The highest-performing raters combined high agreement with consensus, stability across containers, and broad engagement with the corpus—properties that jointly reflect sustained training and accumulated morphological expertise. That the top-performing rater (RI = 0.831) annotated 22 of 31 containers while the lowest-scoring raters tended to contribute to a single container is consistent with the training timeline described in the Methods such that approximately one semester of supervised practice is required before annotators reach intermediate proficiency, and reliability continues to improve with additional experience. The bounded quality weights applied during consensus generation (interval [0.5, 1.5]) reflect a deliberate design choice to modulate rather than exclude lower-performing raters, on the principle that even less experienced annotators contribute information that weighted aggregation can extract. This approach treats annotation quality as a continuous signal to be measured and incorporated rather than a threshold to be enforced consistent with the inseparability of the scientific process and the production of data (3,8,9).

The Context-RF baseline results illuminate a subtle but consequential property of consensus-derived ground truth in that reliability depends on the number of contributing raters. The classifier achieved 50–56% of the majority-vote ceiling on average, establishing that a simple pixel-level model operating on local color and texture features captures roughly half the consensus signal—a gap that represents the contribution of morphological judgment that trained annotators provide and that the consensus workflow is designed to aggregate. Notably, for containers with fewer than seven raters, the Context-RF frequently exceeded the leave-one-out ceiling, a pattern that does not indicate automated superiority over human judgment but rather that the LOO consensus itself is an unstable target when the rater pool is small. In a three-rater container, removing one rater leaves only two, and the resulting majority-vote consensus may not represent the morphological ground truth well enough to serve as a reliable evaluation target. This finding has the practical implication that consensus-derived training labels should be stratified by rater quality and count, and the platform’s Gold/Silver/Bronze tiering system provides the infrastructure to do so. It also suggests that the seven-rater Gold threshold, derived from the convergence analysis, corresponds to a meaningful transition in consensus quality. Specifically, this suggests that below this threshold, consensus estimates are sufficiently unstable that even a simple automated classifier can match or exceed them and that above it the consensus represents a genuine upper bound for automated methods.

Anatolution complements rather than competes with the general-purpose image analysis platforms that have advanced computational histology over the past two decades. CellProfiler provides modular pipelines for a broad range of image analysis tasks and serves a large, diverse user community (27,29,30); Ilastik offers interactive machine learning workflows for pixel and object classification (31); Cellpose provides pre-trained deep learning models for cell detection across tissue types (32,51); and QuPath provides a comprehensive digital pathology workbench with integrated cell detection and annotation tools (52). These tools address algorithmic challenges in segmentation and classification, and they do so effectively—the limiting factor in applying them to comparative neuroanatomy is not their sophistication but the availability of training and evaluation data adequate to the domain. Anatolution addresses this upstream constraint directly, and where existing platforms assume that labeled data already exist or can be produced by individual users, Anatolution provides the infrastructure to produce those labels through a structured, multi-annotator consensus process with integrated reliability assessment. The platform’s outputs are designed to be consumed by any downstream framework—consensus-validated annotations can be exported as image masks compatible with architectures including nnU-Net (36) and Mask R-CNN (43), and because coordinates are stored as resolution-independent percentages, masks can be generated at whatever resolution a given training regime requires. In this sense, Anatolution occupies a distinct position in the analysis pipeline by sitting between raw microscopy images and the segmentation models that existing tools support, filling a gap that is community organizational and methodological rather than algorithmic.

Several limitations of the current platform should be acknowledged for the interested scholar. First, the Nissl-specific segmentation pipeline was optimized empirically on a development set of primate cortical tissue, and its default parameters may not generalize to other species, brain regions, or classical staining methods without adjustment. Second, the pipeline is not designed for fluorescence or electron microscopy, which present fundamentally different image characteristics, and we acknowledge this report’s narrow scope of application. All specimens in the current validation corpus were drawn from previous work (9,44), and while the platform’s architecture is species-agnostic, the generalizability of the segmentation parameters and consensus thresholds to non-primate tissue remains to be established empirically. The consensus workflow, by design, requires multiple trained annotators working on identical image sets—a requirement that makes the platform unsuitable for individual researchers annotating in isolation and that imposes a coordination cost absent from single-annotator tools. Intra-rater reliability, which would provide a complementary measure of annotation consistency within individual annotators over time, was not assessed in this study because the current dataset does not contain re-annotation data. The training investment is substantial, requiring approximately 50 hours of supervised practice before annotators reach intermediate proficiency and with continued improvement over subsequent semesters, this investment is justified by the quality of the resulting data but represents a real barrier to rapid adoption. Perhaps most fundamentally, the value of a consensus annotation platform scales with adoption, turning what is a real problem as outlined above for a single lab into a process for the entire community. The quality of consensus improves as more trained annotators contribute, the utility of exported training data grows as datasets span more species and preparations, and the methodological standards that the workflow enforces become more meaningful as a community of practice forms around them. Anatolution is thus not only a tool but an infrastructure investment whose returns depend on the willingness of the comparative neuroscience community to participate in collaborative annotation—a dependency that is at once the platform’s greatest limitation and, if met, its greatest strength.

Future development priorities follow directly from the limitations described above and from the results of the current validation. Extending the segmentation pipeline to additional classical stains—including hematoxylin and eosin, myelin stains, and silver impregnation methods (22)—would broaden the platform’s applicability beyond Nissl preparations while retaining the same consensus workflow architecture. The multidomain reduced-topography tuning framework described in the segmentation pipeline methods provides a template for this extension. Specifically, the processing chain is stain-general, and adaptation requires primarily the adjustment of color space selection and threshold parameters rather than architectural redesign. Deeper integration with established deep learning frameworks, including nnU-Net (36), Cellpose (32), and Mask R-CNN (43), would streamline the path from consensus-validated annotations to trained segmentation models, reducing the technical friction that currently requires users to write their own preprocessing code.

Completing the downstream model comparison—training nnU-Net models on consensus, single-expert, and single-novice annotations and evaluating all three on a common held-out test set—is the most immediate scientific priority, as it will provide direct evidence for or against the hypothesis that consensus-derived training data produces measurably better segmentation models than individual-annotator labels, and is a next goal. The intra-rater reliability analysis, requiring a dedicated re-annotation protocol, will complement the inter-rater data reported here by characterizing the within-annotator consistency that the Reliability Index currently cannot capture. Moving forward, these and similar analyses will provide a more complete picture of the annotation noise structure that the consensus workflow is designed to manage and leverage.

Anatolution addresses a specific and persistent constraint on quantitative histology: not the absence of segmentation algorithms, but the scarcity of expert-validated training data produced under conditions that make reliability measurable and improvable. The validation results presented here—moderate-to-high inter-rater boundary agreement across 48 raters and five stain types, leave-one-out ceiling Dice values exceeding 0.6 for the best-characterized containers, and 88% polygon coverage by the automated seed detection pipeline—demonstrate that the platform’s consensus-first design produces ground truth annotations whose quality can be measured, weighted, and systematically improved as the annotator community grows. By integrating a stain-specific detection pipeline, a multi-annotator labeling interface, and a consensus workflow with built-in quality control into a single platform, Anatolution provides the comparative neuroscience community with infrastructure for producing the ground truth that its computational methods require. The platform is accessible at https://anatolution.herokuapp.com, with a public demonstration tool available for evaluation without authentication; source code and documentation are available upon request. We invite participation from the community of comparative neuroanatomists, quantitative histologists, and computational biologists whose collective expertise is precisely what consensus annotation is designed to harness.

Building the annotator community is both a practical necessity and a scientific opportunity. Consensus annotation provides a structured pedagogical context in which novice neuroanatomists learn morphological identification under conditions that simultaneously produce useful data—an arrangement in which training and research are not competing demands on time but complementary activities within the same workflow (3,9). As the community grows, the platform’s datasets become more taxonomically diverse, its consensus estimates become more robust, and its training data become more representative of the morphological variation that segmentation models will encounter in practice. The problem of annotation reliability is perhaps most acute for evolutionary neuroscience, where the comparative neuroanatomical lens has revealed extraordinary taxonomic diversity that demands training data spanning the full range of species, brain regions, and staining preparations that the field studies. We thus present Anatolution as part of this special issue on mapping neurobiological diversity, recognizing that the infrastructure for producing high-quality training data is as essential to this enterprise as the algorithms that will consume it.

## Acknowledgement

The authors would like to acknowledge Laura Trice, Iwona Stepniewska, and Huixin Qi for technical support. We also acknowledge the following student users for their role in aggregate data collection over the last decade, in order of contributions made to the datasets presented: Siqi Han, Simran Singh, Makenna Pinkham, Facundo Lodol, Kailee Kaluli, Sooliate Bakare, Eva Baker, Anthony Petrov, Adeyinka Omole, Mustafa Khan, Mia Rode, Emma Lin, Maddy Caliendo, Adedamola Omole, Emily Forsell, Sola Adeyiga, Anya Polonsky, Jared Khan, Susan Lim, Crizaber Cruz, Abigayle Buhay, Hannah Miller, Sophia Vargas, Natalie Marx, Olivia Szelazek, Patrick Kozyra, Bella Peshek, Aanya Sharma, Camille LeBlanc, Daniel Legowski, Nicholas Skowron, Anna Becton, and Georgia Bond.

## Statements

### Statement of Ethics

This study required no protocol or consent to participate as it describes an electronic platform and graphic user interface on the web, and all images were obtained from archived specimens obtained from unrelated research activities primarily related to the Kaas lab who maintained approved study protocols at Vanderbilt University for 50+ years.

### Author Contributions

DJM and JHK designed, funded, and implemented the neuroscientific approach. DJM, ZL, and BG each contributed to the design and construction of the Anatolution project as a toolkit. DJM acquired and managed the image datasets while also collecting annotation data and performing all analyses. ZL and BG were responsible for website design, and BG built the platform. DJM and ZL were both responsible for computational pipeline design and programmatic construction. DJM and ZL were both supported by VIDL fellowships. The project was funded through a VIDL seed macrogrant with Dr. Kaas that also initially funded BG. DJM and BG wrote the manuscript including figures, and then DJM, ZL, and JHK revised the manuscript for intellectual content and merit.

### Data Availability Statement

The authors have made data, code, and documentation available via request or programmatic repository, including an example dataset to show end-to-end reproducibility.

## SUPPLEMENTAL INFORMATION

**SI Figure 1.**
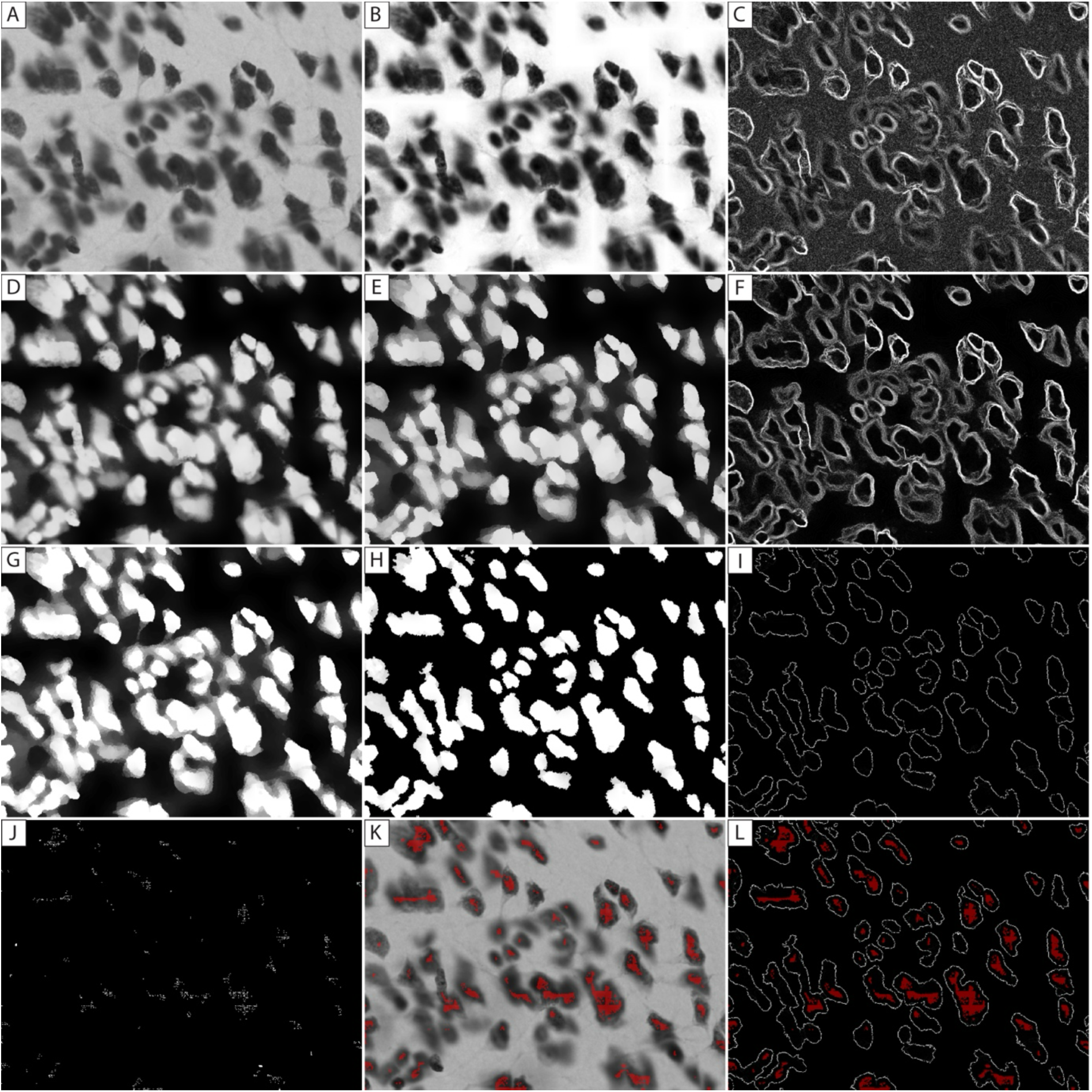
Computational seed detection. Twelve-panel demonstration of the computational seed detection chain and its associated edge diagnostic taps on a single validation tile. The pipeline operates on the original image (A) by extracting the value channel (B) after converting to HSV color space along with calculated edges (C), where cresyl violet absorption contrast is strongest. The protocol then specifies local contrast enhancement (D), and morphological smoothing (E) before calculating edges after smoothing (F). Next, the protocol runs an intensity gate (G), followed by local minima detection (H) and edges on minima (I). The protocol then produces seed locations from the minima (J) that serve as the objective spatial backdrop (K) for the consensus annotations (J) to complete the workflow (see Methods). Scharr edge gradient maps are computed at three diagnostic taps along the chain—after HSV-V extraction (C), after bilateral smoothing (E), and after intensity gating (I) — to visualize how successive processing stages consolidate cell-body boundaries and suppress subcellular texture. Each row of the figure presents two processing stages alongside the edge map computed at the corresponding diagnostic tap, enabling direct comparison of the morphological field and its boundary structure at each stage. All intermediate representations are displayed as 8-bit grayscale images (0–255) unless otherwise noted, and edge maps use percentile-clipped intensity normalization for display. All images depict high resolution cells under x100 objective lens magnification, and the field of view is approximately 80 (y) by 120 (x) micrometers.

**SI Figure 2.**
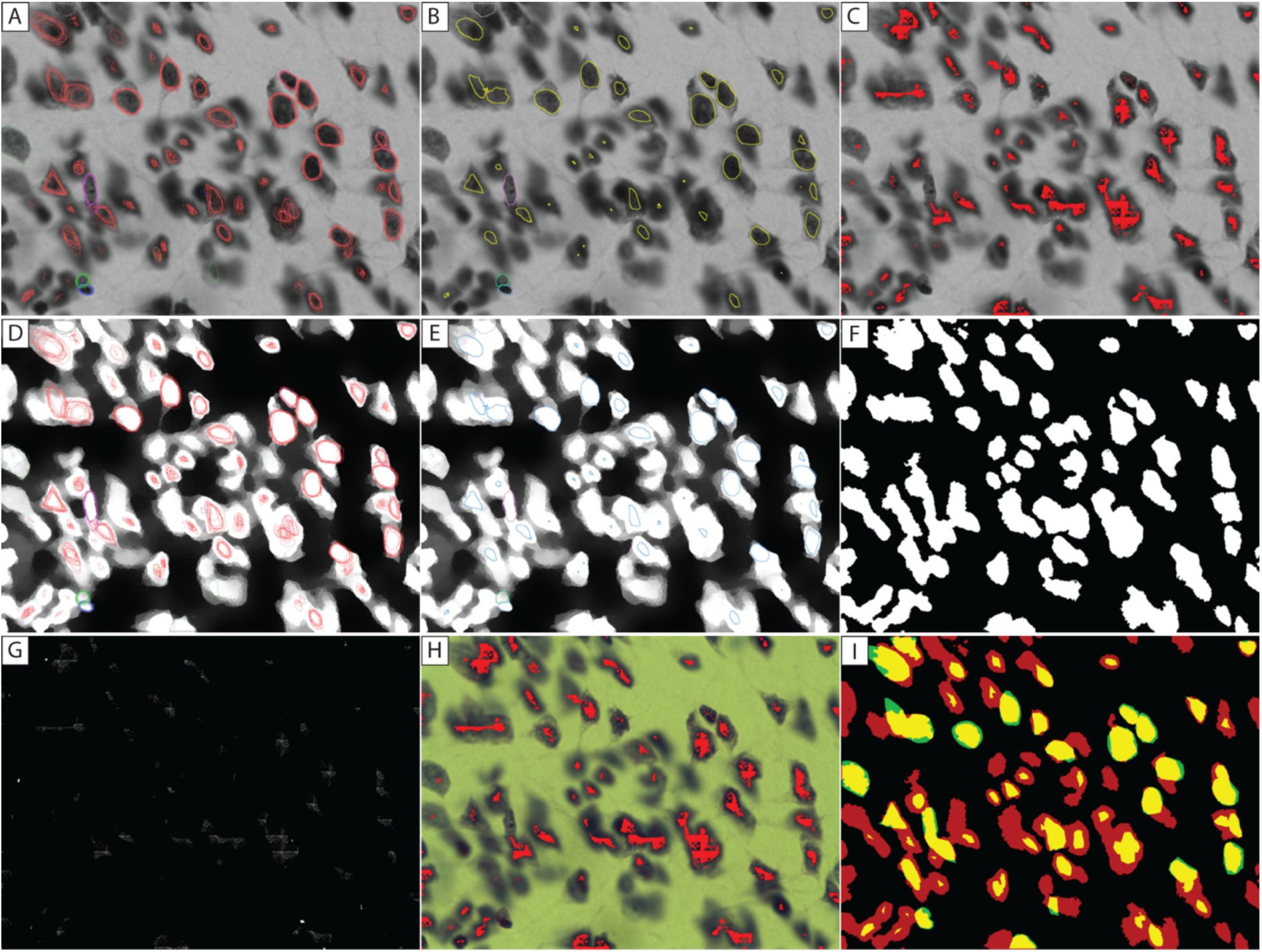
Computational seed validation. Nine-panel validation display for a single image tile integrating consensus annotations (n=12) and computer vision segmentation. This figure integrates the three independent information layers available for each tile—raw tissue morphology, the algorithmically derived reduced-topography scored field, and human consensus annotations—to support visual assessment of pipeline-to-annotation correspondence. The figure is organized as a 3 × 3 grid in which rows correspond to the base image layer (raw luminance grayscale, pre-gate scored field, and diagnostic composites) and columns present progressively more integrated overlays. Panel (A) shows the raw user drawings that are aggregated into a consensus mask (B) and visualized alongside the computer vision seed candidates with calculated edges (C) from the reduced topography (SI Figure 1, and Methods). Panel (D) shows the raw user polygon drawings as well as the consensus mask (E) on the intensity gated segmentation (F). Panel (G) shows the raw seeds in binary that is then converted to red and overlaid on the original color image (H) before also showing the overlay of seeds on consensus (I). Polygon annotation contours are rendered in multicolor overlays (with marker-specific color variants per user visible in Panels A and D). Consensus mask contours appear in yellow on luminance backgrounds and blue on scored-field backgrounds. Seed detections are rendered as binary white on black in the raw data (G), and as red markers for the overlays (C and H). The gated segmentation (F) is converted to red and overlaid with consensus contours (I) to show agreement in yellow, with annotations outside the segmentation depicted in green on black. All images depict high resolution cells under x100 objective lens magnification, and the field of view is approximately 80 (y) by 120 (x) micrometers.

