## Supplemental Figure 1 for "Anatolution, an online platform for consensus morphology"

**SI Figure 1.** Computational seed detection.

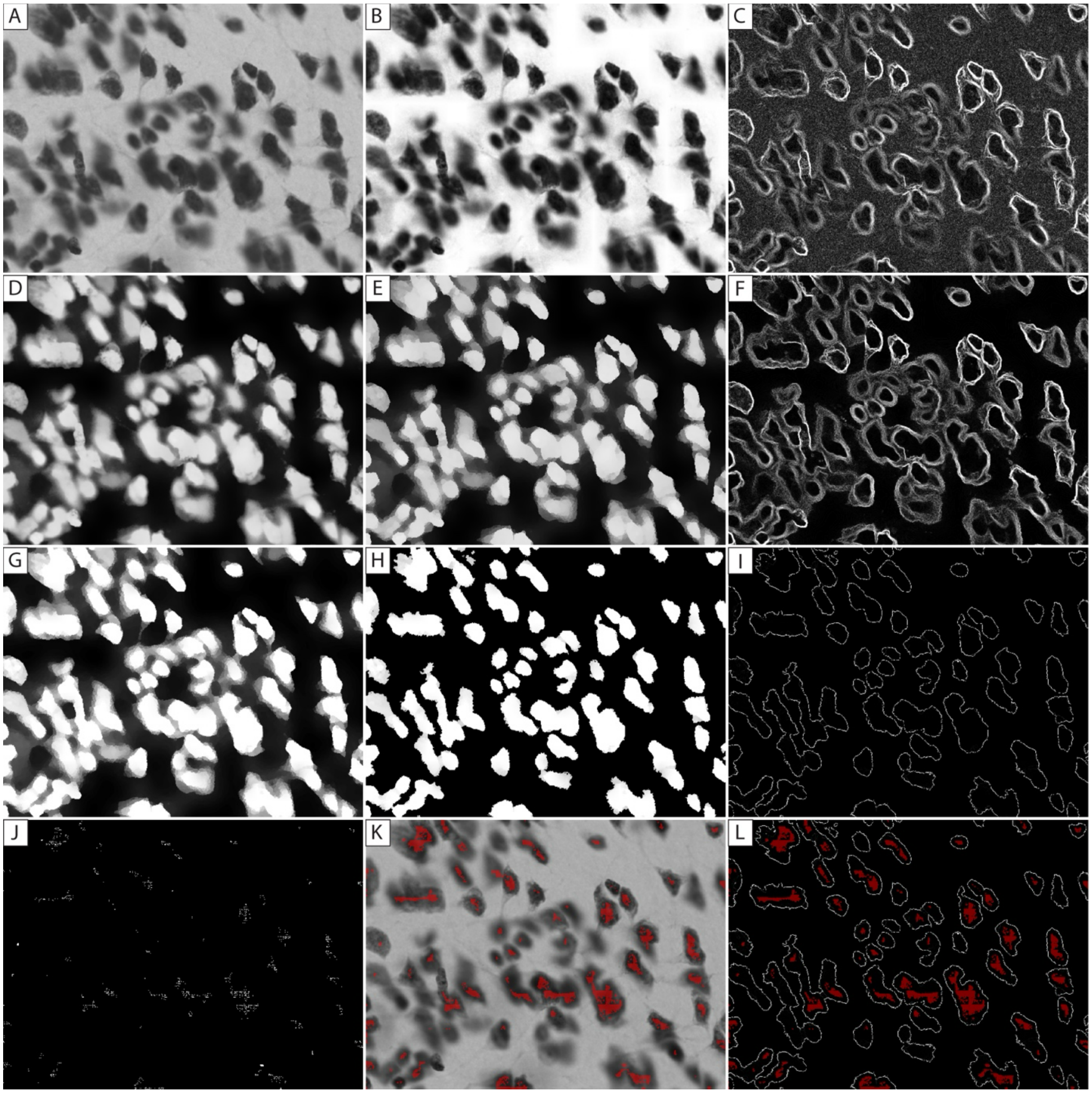

**SI Figure 1 Legend.**

Twelve-panel demonstration of the computational seed detection chain and its associated edge diagnostic taps on a single validation tile. The pipeline operates on the original image (A) by extracting the value channel (B) after converting to HSV color space along with calculated edges (C), where cresyl violet absorption contrast is strongest. The protocol then specifies local contrast enhancement (D), and morphological smoothing (E) before calculating edges after smoothing (F). Next, the protocol runs an intensity gate (G), followed by local minima detection (H) and edges on minima (I). The protocol then produces seed locations from the minima (J) that serve as the objective spatial backdrop (K) for the consensus annotations (J) to complete the workflow (see Methods). Scharr edge gradient maps are computed at three diagnostic taps along the chain—after HSV-V extraction (C), after bilateral smoothing (E), and after intensity gating (I) — to visualize how successive processing stages consolidate cell-body boundaries and suppress subcellular texture. Each row of the figure presents two processing stages alongside the edge map computed at the corresponding diagnostic tap, enabling direct comparison of the morphological field and its boundary structure at each stage. All intermediate representations are displayed as 8-bit grayscale images (0–255) unless otherwise noted, and edge maps use percentile-clipped intensity normalization for display. All images depict high resolution cells under x100 objective lens magnification, and the field of view is approximately 80 (y) by 120 (x) micrometers.
