## Supplemental Figure 2 for "Anatolution, an online platform for consensus morphology"

**SI Figure 2**. Computational seed validation.

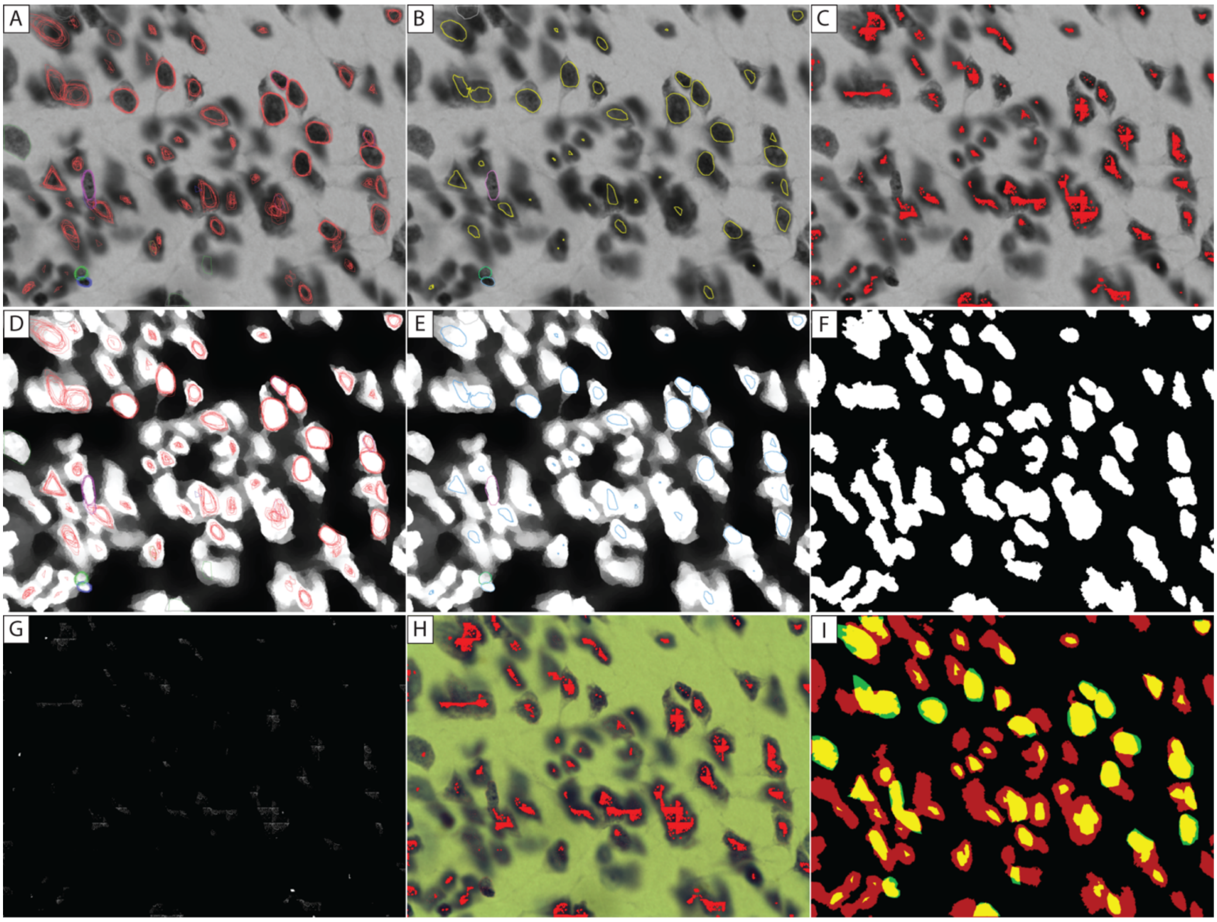

**SI Figure 2 Legend**.

Nine-panel validation display for a single image tile integrating consensus annotations (n=12) and computer vision segmentation. This figure integrates the three independent information layers available for each tile—raw tissue morphology, the algorithmically derived reduced-topography scored field, and human consensus annotations—to support visual assessment of pipeline-to-annotation correspondence. The figure is organized as a 3 × 3 grid in which rows correspond to the base image layer (raw luminance grayscale, pre-gate scored field, and diagnostic composites) and columns present progressively more integrated overlays. Panel (A) shows the raw user drawings that are aggregated into a consensus mask (B) and visualized alongside the computer vision seed candidates with calculated edges (C) from the reduced topography (SI Figure 1, and Methods). Panel (D) shows the raw user polygon drawings as well as the consensus mask (E) on the intensity gated segmentation (F). Panel (G) shows the raw seeds in binary that is then converted to red and overlaid on the original color image (H) before also showing the overlay of seeds on consensus (I). Polygon annotation contours are rendered in multicolor overlays (with marker-specific color variants per user visible in Panels A and D). Consensus mask contours appear in yellow on luminance backgrounds and blue on scored-field backgrounds. Seed detections are rendered as binary white on black in the raw data (G), and as red markers for the overlays (C and H). The gated segmentation (F) is converted to red and overlaid with consensus contours (I) to show agreement in yellow, with annotations outside the segmentation depicted in green on black. All images depict high resolution cells under x100 objective lens magnification, and the field of view is approximately 80 (y) by 120 (x) micrometers.
